# Recurrent interactions can explain the variance in single trial responses

**DOI:** 10.1101/635359

**Authors:** Subhodh Kotekal, Jason N. MacLean

**Affiliations:** Department of Neurobiology, University of Chicago; Committee on Computational Neuroscience

## Abstract

To develop a complete description of sensory encoding, it is necessary to account for trial-to-trial variability in cortical neurons. Using a generalized linear model with terms corresponding to the visual stimulus, mouse running speed, and experimentally measured neuronal correlations, we modeled short term dynamics of L2/3 murine visual cortical neurons to evaluate the relative importance of each factor to neuronal variability within single trials. We find single trial predictions improve most when conditioning on the experimentally measured local correlations in comparison to predictions based on the stimulus or running speed. Specifically, accurate predictions are driven by positively co-varying and synchronously active functional groups of neurons. Including functional groups in the model enhances decoding accuracy of sensory information compared to a model that assumes neuronal independence. Functional groups, in encoding and decoding frameworks, provide an operational definition of Hebbian assemblies in which local correlations largely explain neuronal responses on individual trials.

## 2. Introduction

The earliest single unit recordings in awake primary visual cortex demonstrated that many of the recorded action potentials could not be readily attributed to visual stimulus (Hubel 1959). Across visual cortex, single neuron responses to repeated presentations of a given stimulus exhibit high variability from presentation to presentation (Charles, Park, Weller, Horwitz, & Pillow, 2018; Dean, 1981; Dechery & MacLean, 2018; Goris, Movshon, & Simoncelli, 2014; Heggelund & Albus, 1978; Shadlen & Newsome, 1998; Tolhurst, Movshon, & Dean, 1983). Response variability is largely restricted to neocortex (Scholvinck, Saleem, Benucci, Harris, & Carandini, 2015) and is shared across the neocortical neuronal population (Cohen & Kohn, 2011; Lin, Okun, Carandini, & Harris, 2015). However, neurons from a physiological perspective are capable of being highly reliable (Deweese & Zador, 2004; Mainen & Sejnowski, 1995) suggesting that variance arises primarily from synaptic inputs (Carandini, 2004; Softky & Koch, 1993) and from extraretinal factors that are not visual stimulus such as arousal and locomotion (Goris et al., 2014; Niell & Stryker, 2010).

Neuronal variability can be taken to be purely noise (Faisal, Selen, & Wolpert, 2008), insofar as it is detrimental to the stability of sensory representation. Alternatively, single trial variability may reflect ongoing cortical dynamics associated with sensory processing and representation (Buonomano & Maass, 2009; Harris, 2005). Shared variability may similarly reflect ongoing dynamics (Shimaoka, Steinmetz, Harris, & Carandini, 2019) as pairwise correlations in a population of neurons can affect sensory representation (Abbott & Dayan, 1999; Moreno-Bote et al., 2014; Schneidman, Berry II, Segev, & Bialek, 2006). Neurons are highly interconnected and thus complex network interactions likely shape the activity of neurons (Dechery & MacLean, 2018; Song, Sjöström, Reigl, Nelson, & Chklovskii, 2005). In visual cortex, visually tuned neurons with similar stimulus selectivity are more likely to be synaptically interconnected (Ko, Mrsic-Flogel, & Hofer, 2014) and these connections manifest as specific motifs (Song et al., 2005) which coordinate synaptic integration (Chambers & MacLean, 2016). Moreover, populations of neurons have been shown to exhibit higher-order state correlations (Ohiorhenuan et al., 2010; Schneidman et al., 2006), and topological network features have been shown to shape spike propagation and information transfer (Chambers & MacLean, 2016; Hu et al., 2018; Womelsdorf, Valiante, Sahin, Miller, & Tiesinga, 2014). However, the relative role of local synaptic connectivity, as compared to stimulus-related input or more global variables such as locomotion, in the generation of a sensory representation, particularly in real time, remains unclear. A comprehensive theory of primary visual cortex must encompass local correlations instead of treating neurons as independent units.

The cell assembly hypothesis, laid out by Donald Hebb, proposes that neurons are organized into mutually excitable groups called “assemblies” which are strongly coactive (Harris, 2005; Hebb, 1949). Under this hypothesis, sensory representations and cortical state dynamics manifest as sequences of assembly activations and internal state dynamics need not be deterministically tied to stimulus from trial-to-trial (Buonomano & Maass, 2009; Harris, 2005; Hebb, 1949; Scholvinck et al., 2015). Importantly, the assembly hypothesis suggests that single trial dynamics are strongly influenced by the group of coactive neurons rather than entirely by the external stimulus.

In this work, we investigate whether the variance in V1 neuronal activity over the time course of single trials is best explained by visual stimuli, locomotion, or by coactive groups of V1 neurons. We define coactive groups of neurons, which we term as “functional groups”, by local pairwise correlations of activity within short time intervals after accounting for stimulus and running effects. We used a generalized linear model (GLM) on calcium imaging data of visually-evoked activity recorded from V1 in freely running mice. In the model, we relate the target neuron’s activity during single trials to the functional group’s activity, stimulus condition, and mouse running speed. Functional group coactivity is understood to be informative of a target neuron’s activity if inclusion of coupling in the GLM improves predictions of the target’s neuron activity, which is suggestive of the importance of internal circuit dynamics in shaping single trials (Harris, 2005; Harris, Csicsvari, Hirase, Dragoi, & Buzsáki, 2003). With the functional group as an operational definition of an assembly, we provide concrete descriptions of the timescales, numerical scale, correlation structure, and computational capabilities of assemblies.

## 3. Results

Trial-to-trial, the activity of individual neurons in visual cortex is highly variable despite ostensibly identical conditions (Shadlen & Newsome, 1998). To evaluate local and global effects on single trial variability, we built an encoding model using a generalized linear model (GLM), comprised of three terms: neuron coupling, stimulus and running speed. To determine each term’s contribution, we constructed multiple restricted GLMs. Each excluded a term of interest and we compared test set predictions to those of the unrestricted GLM (i.e. the model containing all three terms). Relative change in a restricted model’s test set performance indicated the excluded term’s importance in modelling single trial activity. For example, a large increase in a restricted model’s test set error indicated that the excluded variable was informative to single trial activity. A marginal change indicated that the excluded variable was uninformative.

Notably, we used prediction performance rather than the GLM coefficients to determine the importance of model terms. The full set of variables that truly explain V1 single trial activity is not *a priori* known, nor experimentally observable. Coefficient estimates would be different if the omitted variables were included in the model, and may lead to erroneous conclusions; this is omitted variable bias (Stevenson, 2018). Consequently, we used the unrestricted GLM’s mean squared error (MSE) on the test set data as a benchmark against which we compared each restricted model, and thus evaluated the importance each variable.

### 3.1. Mouse Visual Cortical Data

The imaging data used throughout this manuscript was described in a previous study (Dechery and MacLean 2018). Briefly, the activity of L2/3 excitatory neurons expressing GCaMP6s in mouse visual cortex (73–347 neurons; n = 8 animals; 21 distinct fields of view; Fig. 1A) were imaged using two photon laser scanning microscopy (25–33 Hz; Dechery and MacLean 2018; Sadovsky et al 2011). Mice were awake and allowed to freely run on a linear treadmill while viewing drifting square-wave gratings presented in 12 directions in pseudorandom order interleaved with mean-luminance matched grey screen; each trial was a 5-minute block of stimulus presentation in this format.

**Fig. 1:**
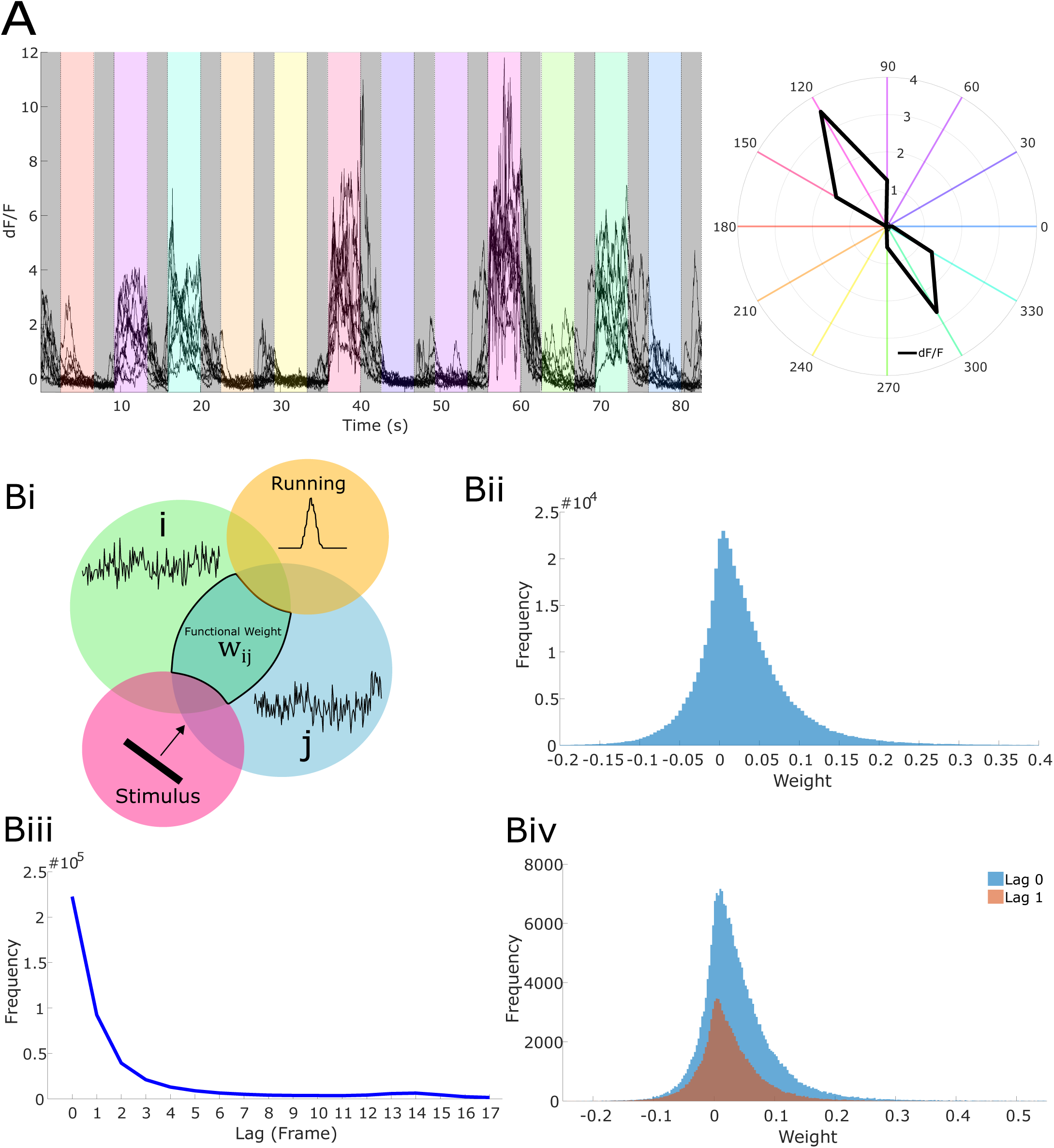
(A) Left, anecdotal example fluorescence traces of neuronal responses to oriented drifting gratings (neuron 260, dataset 3). Right, polar plot of the average response of anecdotal neuron averaged across time bins and stimulus presentations. Colors in both panels correspond to different grating drift directions. (Bi) Diagram of partial correlation measure used for coupling in generalized linear models (GLMs); the partial correlation between neurons *i* and *j* is the correlation in the activities after accounting for stimulus and running induced coactivity. In the diagram, the partial correlation is denoted and outlined as the region of intersection between the regions of neurons *i* and *j*, but not including the regions of stimulus and running. (Bii) Functional weight (i.e. partial correlation magnitude) distribution across population. (Biii) Functional lag distribution with respect to lag length across population. (Biv) Functional weight distribution segregated by lag 0 and lag 1.

### 3.2. Experimentally measuring coupling coefficients for use in a GLM

Visual stimuli (Dechery & MacLean, 2018) were repeatedly presented in identical 5-minute blocks. Partial correlations, i.e. *functional weights*, were computed for each pair of neurons (Fig. 1B), where each functional weight captured the reliability of coactivity after accounting for shared changes in activity due to stimulus and running speed. This measure is analogous to “noise correlations” (Cohen & Kohn, 2011), but also allowed us to account for the running-induced changes in circuit activity (Niell & Stryker, 2010). To summarize the temporal component of coactivity, we computed the mean traces of each neuron and constructed a cross-correlogram, the maximum value of which denoted the *functional lag* between the neurons. Functional connections were often not symmetric, and the direction of the lag gives a direction of the connection. The associated functional lags most frequently were 0 frames (Fig. 1Biii) and frequency decreased rapidly with lag amplitude; most coupling was bidirectional (lag 0) (mean = 50.1%, sem = 2.8%, mean over n = 21 datasets). For each neuron, the set of other neurons with functional connections directed towards the given neuron was called its *functional group*. The average number of incoming functional connections for a target neuron (i.e. functional group size) was 43.18 ± 3.76 (mean ± s.e.m.). We employed these experimentally measured functional connections as coupling coefficients in a GLM rather than fitted coefficients to mitigate omitted variable bias (Stevenson, 2018).

#### 3.2.1 Performance of the unrestricted GLM

To assess the accuracy of a simple GLM model (consisting of coupling coefficients, a stimulus term, and a running term), we evaluated our ability to predict single trial neuron fluorescence. To model the stimulus, we first determined neuronal response properties by computing the average stimulus-dependent response (averaged across all stimulus presentation time bins and all blocks). Neurons significantly tuned to orientation or direction were labeled as *tuned* with the procedure described in (Dechery & MacLean, 2018); all others were labeled *untuned*. For tuned and untuned neurons, the stimulus term in our GLM was the experimentally observed average stimulus-dependent response and overall average response respectively. The running term for both types of neurons was given by the mouse’s running speed as measured by a rotary encoder attached to the axle of the mouse treadmill. Since most functional weights (70.62% ± 2.76%) (mean ± s.e.m where mean is over datasets) had associated lags with values 0 or 1 (in units of roughly 30 *ms* time bins), we only included these coupling lags in our model. We note that each lag 0 connection was counted twice to indicate bidirectionality. Pooling across neurons, the unrestricted GLM exhibited accurate predictions of single trial neuronal fluorescence (fluorescence traces normalized for each neuron) as indicated by a small test set mean squared error (MSE) (median = 0.0402, IQR = 0.0616, [25^th^ percentile = 0.0233, 75^th^ percentile = 0.0849] Fig. 2A). The test set MSE was stable over the entirety of stimulus presentation since the first second after stimulus onset exhibited similar test MSE relative to the entire trial (median = −0.80%, IQR = 17.54%).

**Fig. 2:**
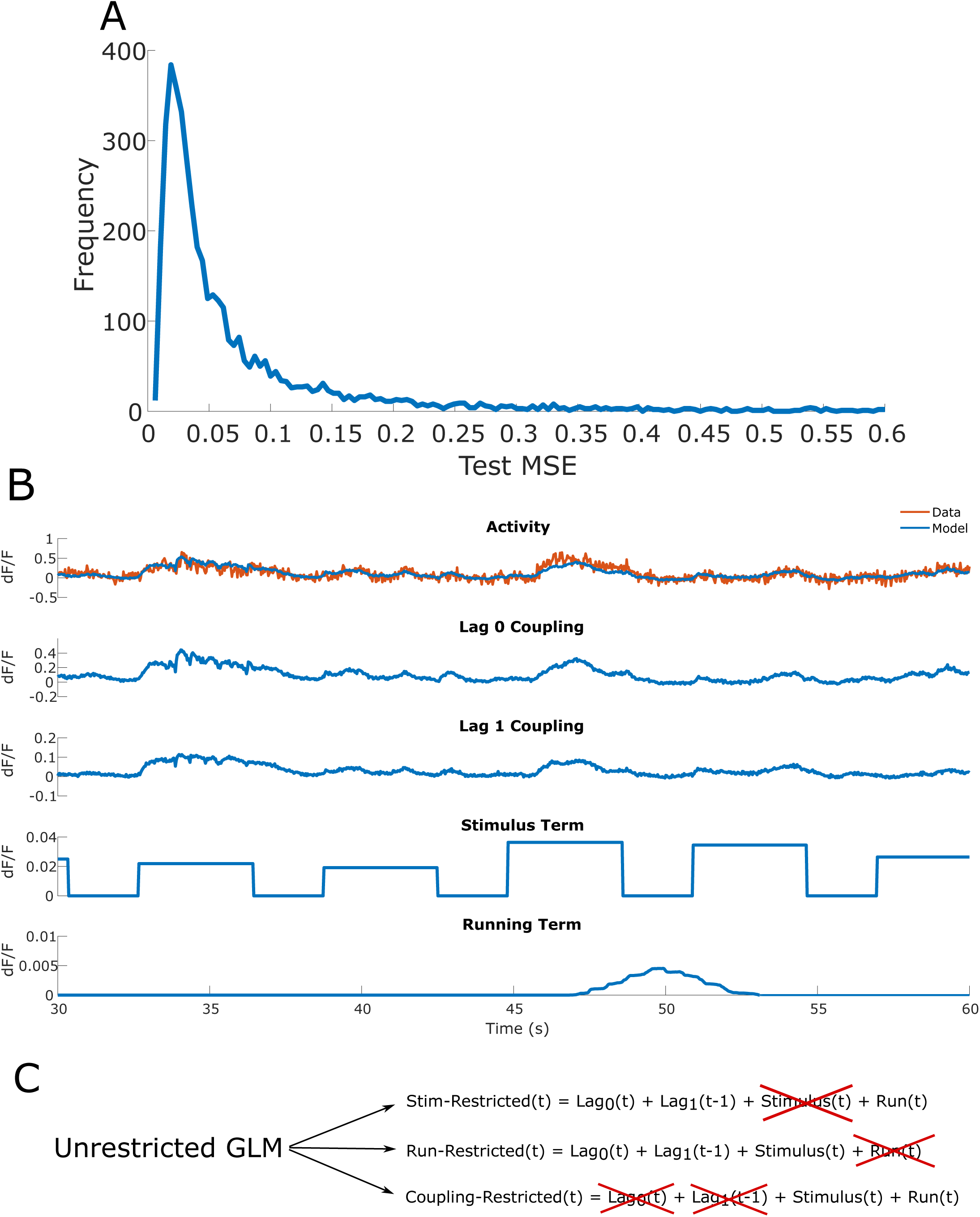
(A) Distribution of the percent change in test set mean-squared error (MSE) for the unrestricted GLM across all neurons. The unrestricted GLM contains all three model terms, which are the coupling coefficients, stimulus term (average stimulus-dependent response), and running term (running speed given by rotary encoder). Across stimulus/grey conditions, the unrestricted GLM models the single trial responses of many neurons accurately with a right skewed distribution. (B) Top, time-varying model terms and subsequent prediction for anecdotal neuron (neuron 125, dataset 3). Bottom, diagram for construction of restricted models. The relative changes in test set MSE of the restricted GLMs compared to the unrestricted GLM indicate the importance of the excluded term in generating accurate predictions of single trial responses. For example, if the stimulus-restricted GLM exhibits a minimal test set MSE increase relative to the test set MSE of the unrestricted GLM, then the stimulus is understood to be uninformative to predicting single trial responses.

The accurate predictions of the unrestricted GLM allowed us to next evaluate which individual terms best contribute to capturing single trial neuronal dynamics. To isolate the relative contribution of each term to the prediction accuracy of single trial fluorescence, we excluded the stimulus or running term (*stim-restricted GLM* and *run-restricted GLM* respectively) and compared these models against the prediction accuracy of the *unrestricted GLM* (Fig. 2C). Each of the three GLMs were individually fit on the same training data and compared on the same test data (roughly a 70/30 data split). Additionally, we fit GLMs in which all neuronal couplings were excluded (i.e. *coupling-restricted GLM*) to determine the extent to which the functional group contributed to accurate prediction of single trial responses.

#### 3.2.2 Performance of the stimulus-restricted GLM

We modeled the stimulus in two ways to examine the extent to which averaged summaries of stimulus-dependent responses can contribute to accurate predictions.

First, as in the unrestricted GLM above, we chose to model the visual stimuli as trial averaged responses given that the transform(s) of visual information as it transits from retina to LGN to layer 4 to layer 2/3 is not fully known. In particular, we averaged the empirically measured stimulus-dependent fluorescence across all time bins per stimulus condition across all trials (each trial being a single 5-minute block) (Fig. 3A), as described above. We then fit a GLM containing all terms (i.e. unrestricted GLM) and a GLM with all terms except the stimulus term (stimulus-restricted GLM). When comparing between our two models of the stimulus, we restricted our analysis of the results to only the drifting grating (i.e. non-grey stimulus) imaging frames.

**Fig. 3:**
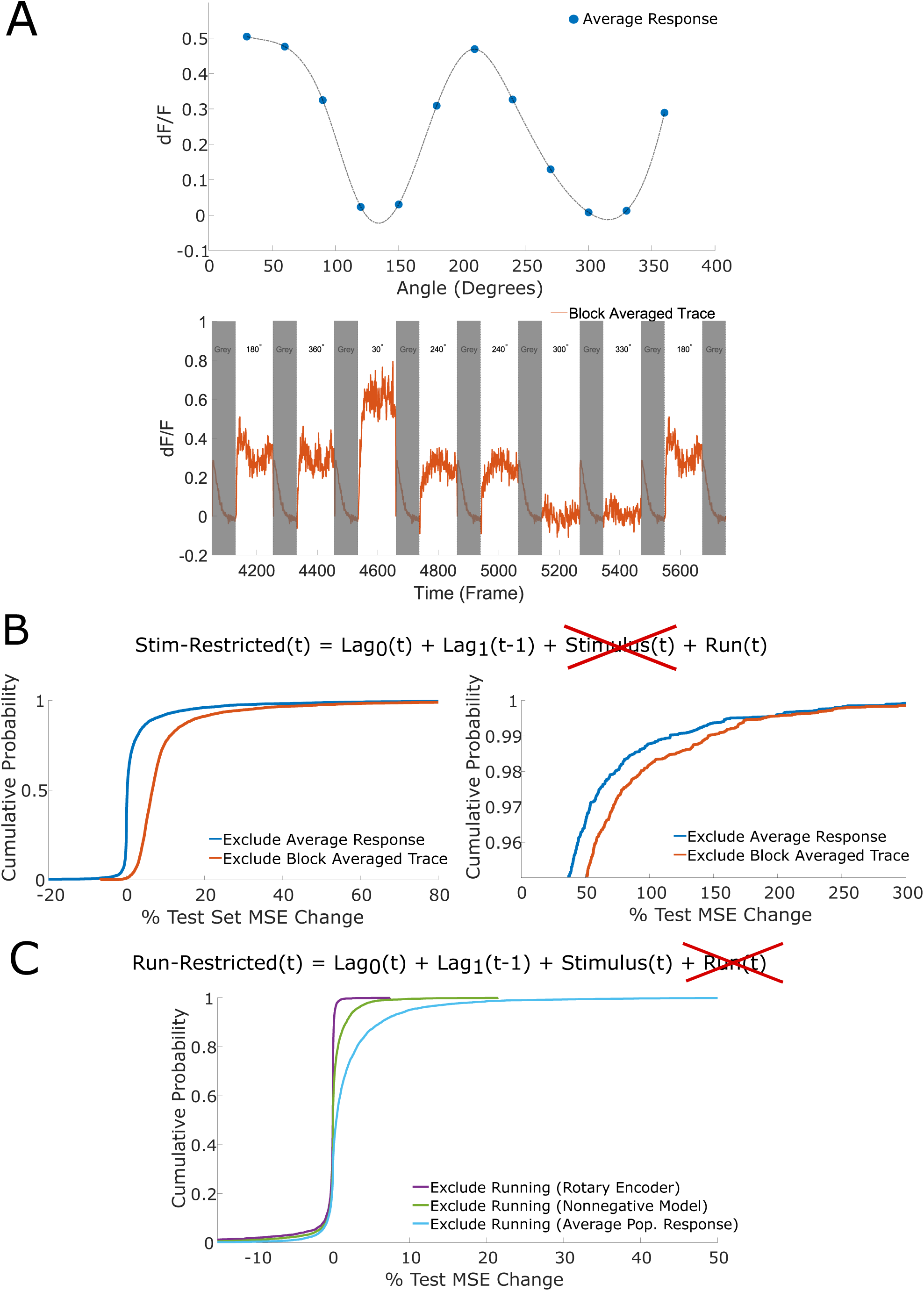
(A) Top, stimulus-dependent average response profile for an anecdotal example neuron (neuron 2, dataset 3). The average response is a grand mean, i.e. the average response of a neuron to a given stimulus is the neuron’s response to the stimulus averaged across all time bins of all presentations of the given stimulus. In effect, the average response is the average fluorescence change the neuron exhibits in response to a given stimulus. Bottom, an example of a block averaged trace for the same neuron (neuron 2, dataset 3). The block averaged trace of a neuron for a given stimulus is the trace obtained when averaging traces across all blocks (i.e. trials) and all presentations of a given stimulus within each block. In effect, the block averaged trace represents the average trace of fluorescence changes in response to a given stimulus. For the GLMs using the block averaged trace model, we restricted our analysis only to the stimulus frames and excluded the grey frames which are obscured here using grey bars. (B) Left, cumulative distribution functions of the percent change of test set MSE across stimulus frames for the stimulus-restricted GLM using the average response stimulus model and the stimulus-restricted GLM using the block averaged trace stimulus model. Right, zoom in on 95^th^ percentile. (C) Cumulative distribution functions of the percent change of test set MSE across all frames for the run-restricted GLMs using the rotary encoder running model, nonnegative model coefficients, and average population response running model.

Pooling across neurons, the stimulus-restricted GLM exhibited marginal test set MSE increase (all frames: median = +0.20%, IQR = 1.43%, stimulus frames: median = +0.22%, IQR = 1.79%, Fig. 3B). The stim-restricted GLM’s marginal MSE increase suggests that knowledge of the average response does not improve single trial predictions when conditioning on the functional group’s activity and the running speed. It must be noted, however, that neurons in the 95^th^ percentile of stimulus-restricted GLM test set MSE increase exhibit a large increase (median = +29.73%, IQR = 29.11% across stimulus frames, Fig. 3B), indicating that average response captured a large component of the single trial dynamics for a small subset of neurons.

Our second model of the stimulus attempted to account for the temporal dynamics of a stimulus-dependent response. In this model, tuned and untuned neurons were treated the same. For each neuron, we averaged its fluorescence trace in response to a given stimulus across all presentations and across all trials to obtain the “block averaged trace” for a given stimulus (Fig. 3A). The block averaged trace was thus obtained after averaging over 9-15 traces depending on the dataset; as noted above, we tested this model only on stimulus frames (i.e. non-grey frames). In order to avoid averaging over different numbers of presentations, we took a 50/50 split of the data for training and testing the models. Notably, by considering fewer trails, we necessarily were less able to completely isolate and account for the other two variables, coupling and running speed using partial correlation. As before, we fit an unrestricted GLM and the stimulus-restricted GLM, and we restricted analysis to the drifting grating (i.e. non-grey stimulus) frames.

Across the population, this variant of the stimulus-restricted GLM again exhibited marginal increase in test set MSE (median = +6.64%, IQR = 5.02% across stimulus frames, Fig. 3B). For the majority of neurons, averaged summaries of stimulus-dependent responses are ill-equipped to explain single trial activity even when accounting for the average temporal component of dynamics. Again, neurons in the 95^th^ percentile of stimulus-restricted test MSE increase exhibited large MSE increase (median = +50.59%, IQR = 36.87% across stimulus frames, Fig. 3B). The remainder of our analysis in this paper used the average response model for the stimulus term unless otherwise specified.

#### 3.2.3 Performance of the run-restricted GLM

We evaluated the contribution of the running term (running speed measured by rotary encoder) to accurate predictions of single neuron fluorescence transients. The run-restricted GLM exhibited marginal changes in test set MSE compared to the unrestricted GLM (median = −0.01%, IQR = 0.20%, Fig. 3C), indicating that inclusion of running speed failed to improve prediction accuracy when conditioning on functional group activity and the average response. In numerous cases, we obtained a negative coefficient for the running term as we did not constrain model coefficients to be nonnegative. While we do not focus on the coefficients in the majority of the paper, it was necessary to address these particular values because it is well known that running speed strongly modulates spike rates (Niell & Stryker, 2010), and our data similarly showed enhanced population activity during periods of running (Dechery & MacLean, 2018). Given that (1) we are inevitably subsampling visual cortical circuits and thus cannot include all relevant variables in the GLM, and (2) we had found that run speed coefficients are mostly small in magnitude (median = −0.0001, IQR = 0.0062), it was possible that sign errors were due to omitted variable bias (Stevenson, 2018). Consequently, we refit the unrestricted GLM and the run restricted GLM with the additional constraint that coefficients of all model terms (i.e. coupling, stimulus, and running terms) be nonnegative. Again the change in test set mean squared error (MSE) relative to the unrestricted GLM was small (median = 0.0060%, IQR = 0.49%, Fig. 3C). Although we observed global increases in neuronal activity across all imaged neurons during running, the inclusion of a running term did not substantially contribute to accurate prediction of neuronal fluorescence in held out data.

To account for the possibility that running speed does not linearly drive population activity, we constructed a second running model. We used the time-varying average population response as the running term since running is known to globally modulate the population. Fitting and testing the corresponding unrestricted and run-restricted GLMs, we found that the average population response running model captured more of the dynamics than the nonnegative rotary encoder model, but was still relatively uninformative as changes in test set MSE were small (median = 0.45%, IQR = 2.32%, Fig. 3C). In the remainder of our analysis, we used the rotary encoder running model when not otherwise specified.

#### 3.2.4 Performance of the coupling-restricted GLM

The above results suggested that inclusion of the stimulus condition and running speed were largely uninformative to generating accurate predictions of single trial neuronal dynamics in the full GLM. However, it was unclear whether these terms enable accurate predictions when *not* conditioning on functional group activity. We fit a GLM with all terms (with the average response stimulus model and rotary encoder running model) except coupling on the training set. The coupling-restricted GLM exhibited a large test set MSE increase relative to the unrestricted GLM (median = 28.93%, IQR = 47.03%, Fig. 5A top right; across all frames), demonstrating that the functional group captured variance unexplainable by the average stimulus response and running speed. This result held when the block-averaged trace was used as the stimulus term (with the rotary encoder running model), with the coupling-restricted GLM exhibiting a large increase in test set MSE (median = 22.35%, IQR = 39.80%; across stimulus frames). Collectively these results indicate that the couplings between neurons contribute substantially to accurate model predictions.

### 3.3. GLM prediction accuracy is sensitive to the specifics of topology and large weights

While it was clear that the functional group coupling enabled accurate predictions of single trial activity, the importance of specific topological features of the functional group was unclear. Is prediction more sensitive to the precise weight values or the underlying presence of specific connections? To explore these relationships, we employed graph theoretic methods to manipulate functional group weights and connections. In this framework, a connection is referred to as an edge.

Previous work has shown that weak functional weights are uninformative of single trial dynamics (Dechery & MacLean, 2018). To examine the importance of specific topological features amongst the strongest weighted edges in the functional group, we first identified the edges with weights in the top quartile of magnitudes (termed the “strong edges” and “strong weights”) and permuted the weights (along with the corresponding lag) among these edges. This permutation scheme preserved the topology of the functional groups but shuffled the associated weights (Fig. 4A). The new weights of the functional group were then substituted into the unrestricted GLM. We did not refit the unrestricted GLM, but rather computed the change of MSE on the training data (here, we revert to analyzing all frames). Refitting the GLM would have optimized model coefficients to minimize model error, thus obscuring the effect of shuffled weights. This shuffling procedure was repeated 1000 times, and resulted in a large median increase of training MSE (median of median increase = +23.57%, IQR of median increase = 57.99%, Fig. 4B top). These results demonstrate that the predictive abilities of functional groups are highly sensitive to the precise value and association of each strong functional weight to a specific edge.

**Fig. 4:**
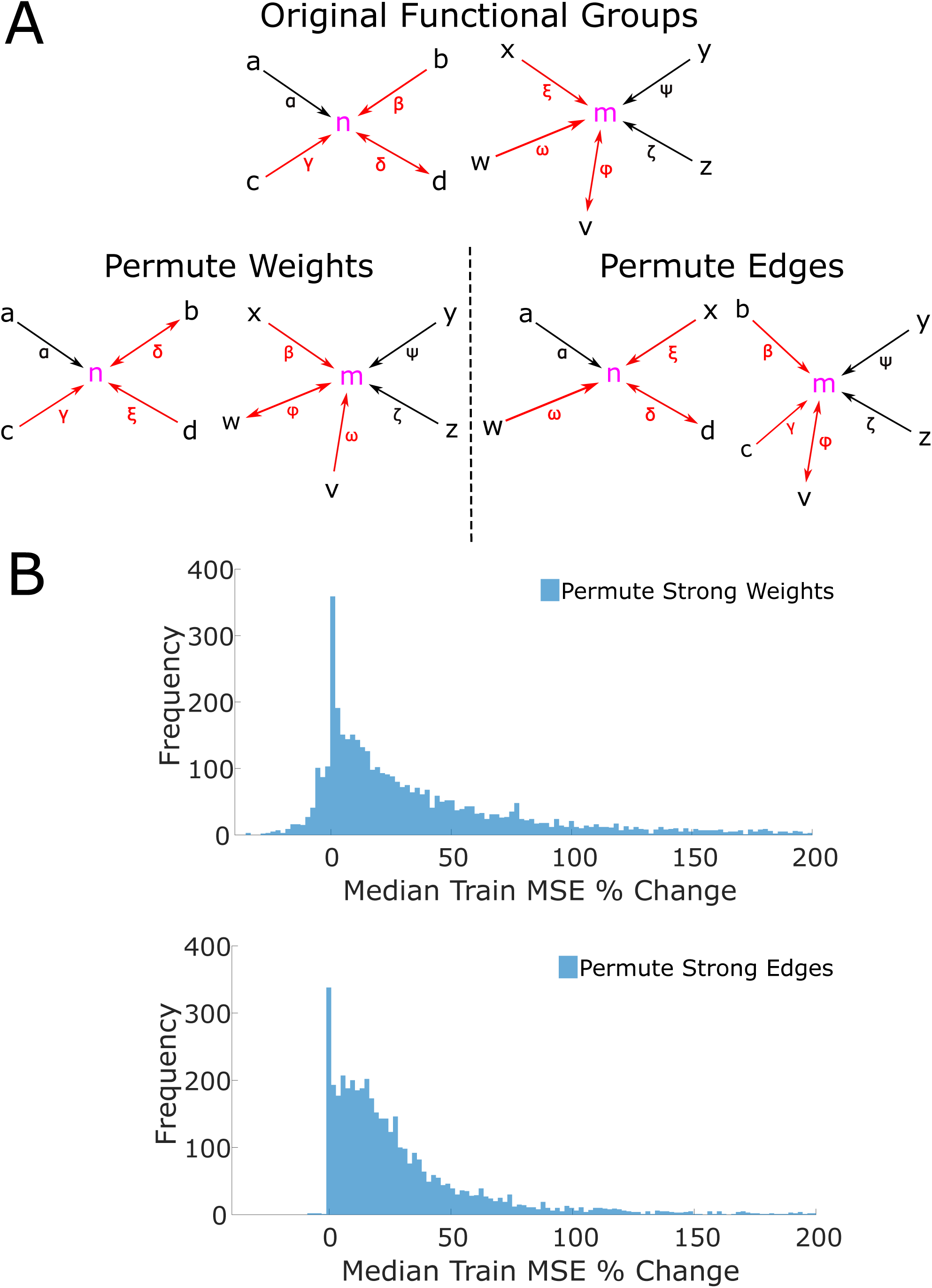
(A) Diagram of permutation methods. Top, example of the functional groups for neurons n and m. The roman letters denote neurons that are coupled to the target neurons. Greek letters denote the weight of the edge (i.e. the functional weight). The arrows show the directionality of the coupling, with bidirectional coupling indicating a lag 0 edge. Red coloring of edges and weights indicate that the edge is “strong” (i.e. has an edge in the top quartile of magnitudes). Only strong edges and weights are permuted in the two shuffle methods. Bottom left, an example permutation of the strong weights of the original functional groups. Note that each functional group retains the neurons in its group, but the strong weights (denoted by the red Greek letters) are freely permuted, even between functional groups. Note that the lag is permuted along the weight, and so the directionality of the coupling follows the weight when it is permuted. For example, note that the weights β, δ, and ξ are all permuted, and the bidirectional coupling associated with δ follows the permuted weight δ. Bottom right, an example permutation of strong edges of the functional groups. When permuting edges, the neuron memberships of the functional groups change as strong edges are permuted in and out of each functional group. For example, neurons *b* and *c* are permuted into the functional group of m from the functional group of *n*. Similarly, neurons *x* and *w* are permuted into the functional group of n from the functional group of n. By permuting edges, new functional group topologies (in the sense of neuron membership) are instantiated. (B) Top, distribution of the median percent change of training set MSE across all 1000 permutations of strong weights. Bottom, distribution of the median percent change of training set MSE across all 1000 permutations of strong edges. These distributions indicate that accurate predictions from the model are sensitive to perturbations of both strong weights and edges of functional groups.

Second, we again identified the edges with strong weights (termed “strong edges”). Now, rather than shuffle only the weights, we shuffled the strongest edges and thereby generated new topologies (Fig. 4A). This allowed us to determine the sensitivity of prediction ability to the underlying strong edges. To elaborate, this shuffling procedure shuffled *all* of the strong edges across *all* functional groups. After shuffling, each neuron’s functional group was changed since the composition of a functional group is defined by the specific incoming edges that each neuron receives. Again the GLM was not refit for the same reasons articulated above, and this procedure was repeated 1000 times. Shuffled edges resulted in a large median increase of training set MSE (median of median increase = +20.00%, median of IQR = 31.43%, Fig. 4B right).

Thus, both the specific weights and the specific connections in the upper quartile of edges define the functional group which in turn underlies the impressive performance of the coupled model. Large weights thus explain the importance of specific edges in the functional group for accurate predictions and are a signature of the important timescales and weights (regardless of sign) of the functional group; consequently, the timescale and weights of Hebbian assemblies can be further characterized by observing which timescales and weight sign correspond to the edges with large weights in the functional group.

### 3.4. Recurrent coupling coefficients contribute to accurate predictions

Given the importance of the functional group, we next asked how its temporal structure related to prediction. To do so we constructed two models, GLM_0_ and GLM_1_, which respectively contained only a lag 0 or a lag 1 coupling term, and compared prediction performance to an unrestricted GLM. We additionally examined the large weight distributions for lag 0 and lag 1 coupling.

We found that the GLM_0_ exhibited a marginally increased test set MSE while GLM_1_ exhibited a large test set MSE increase as compared to the unrestricted model that contained both coupling terms (GLM_0_ median = +0.63%, IQR = 2.16%, GLM_1_ median = +9.81%, IQR = 21.10%, Fig. 5B). This result demonstrates that lag 0 coupling is necessary to generate accurate single trial predictions and suggests that that prediction-relevant coactivity between neuronal pairs occurs within a window less than or equal to ∼30 *ms*. Hence, the predictive ability of functional group coactivity is driven by the functional group’s bidirectional edges.

**Fig. 5:**
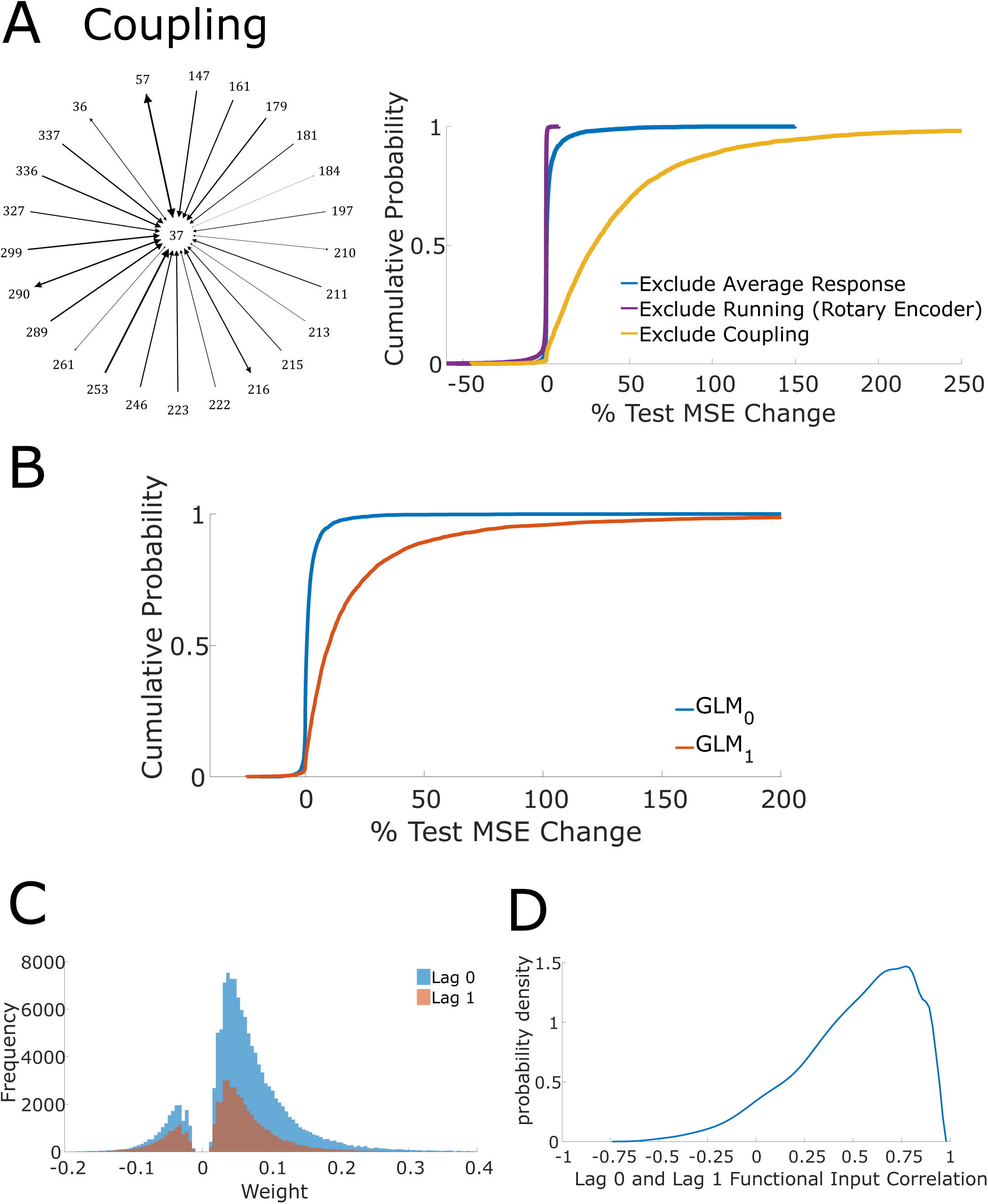
(A) Left, example of functional group of an anecdotal neuron (neuron 37, dataset 3). Diagram shows all neurons with nonzero weight (across all lags) that have directed coupling outgoing to neuron 37 in dataset 3; these neurons constitute the functional group for neuron 37 in dataset 3. Directionality is indicated by arrows and edge weight is denoted by thickness of the edge and arrow head size, with thickness being a monotonically increasing function of weight. Top right, cumulative distribution functions for the percent change in test set MSE across all frames for stimulus-restricted, run-restricted, and coupling-restricted GLMs using the average response stimulus model and the rotary encoder running model. (B) Cumulative distribution function of percent change in test set MSE of GLM_0_ and GLM_1_ with respect to the unrestricted GLM. The exclusion of lag 0 results in a large increase in test set MSE, indicating that lag 0 edges are informative to predicting single trial responses. The exclusion of lag 1 results in a marginal increases of test set MSE, indicating that lag 1 edges are mostly uninformative. (C) Strong weight distribution segregated by lag 0 and lag 1 (“strong” weight means functional weight in top quartile of magnitudes). Strong edges are more likely to be lag 0 edges, indicating why lag 0 edges are informative to accurate predictions. (D) Probability density estimate of Pearson correlation between lag 0 and lag 1 coupling across all neurons.

The sufficiency of lag 0 weights and insufficiency of lag 1 weights to predict single trials is informed by the prevalence of strong weights associated with lag 0 edges (“strong” defined as belonging to the top quartile) (Fig. 5C). Furthermore, we observe that strong coupling exhibits more lag 0 weights than lag 1, which is consistent with our previous results regarding large weights. While it is the case that lag 0 and lag 1 functional input are linearly correlated (median r = 0.56, IQR = 0.40, Fig. 5D), collinearity cannot totally explain the sufficiency of lag 0. If activity from lag 0 and lag 1 coupling are collinear, then GLM_1_ should have performed comparably to GLM_0_, as the lag 1 coupling GLM coefficient would be modified linearly to account for the absence of lag 0 coupling; yet, GLM_1_ performed worse than GLM_0_.

### 3.5. Necessity of positive coupling and redundancy of negative coupling

We next asked to what extent prediction was attributable to positive or negative coupling coefficients. Negative correlations have been shown theoretically to increase coding capacity (Sompolinsky, Yoon, Kang, & Shamir, 2001) and consequently were of particular interest here. Two models, GLM^+^ and GLM^-^, were constructed where only the positive weight term or negative weight term was included respectively; additionally, an unrestricted GLM was fit. Stimulus and running speed terms were included in all models; however, we did not restrict the set of lags under consideration. GLM^+^ exhibited a marginal increase in test set MSE (median = +0.24%, IQR = 1.43%, Fig. 6A) whereas GLM^-^ exhibited a large increase in test set MSE (median = +25.38%, IQR = 36.37%, Fig. 6A). These results indicate that positive weights are sufficient for accurate prediction while negatively correlated neurons contributed minimally. This result further implies that the positive and negative coupling model terms are not collinear as GLM^+^ and GLM^-^ did not exhibit comparable performance. The performance difference of GLM^+^ and GLM^-^ is notably marked by the prevalence of strong positive as compared to negative weights. Across datasets, the top quartiles of weight magnitudes are comprised of many positive weights and few negative weights (Fig. 6B).

**Fig. 6:**
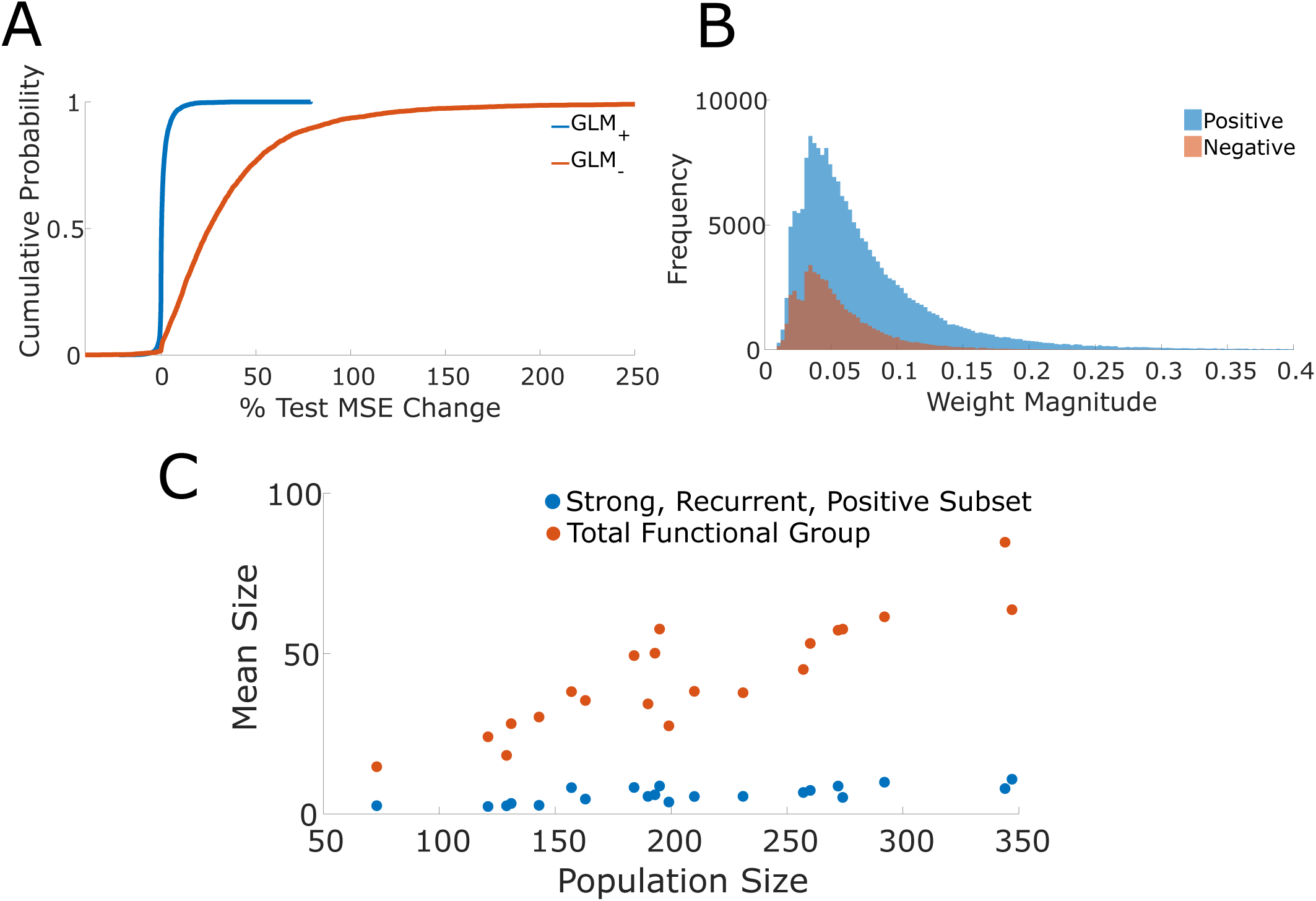
(A) Cumulative distribution function of percent change in test set MSE of GLM_+_ and GLM_-_ with respect to the unrestricted GLM. The exclusion of edges with positive weights results in a large increase of test set MSE while the exclusion of edges with negative weights results in only minimal increases. These results suggest that edges with positive weights are informative while edges with negative weights are uninformative in generating accurate predictions of single trial responses. (B) Distribution of strong weight magnitudes segregated by positive and negative weights. Strong weights are more likely to be positive rather than negative, thus indicating why positive weighted edges are informative to accurate predictions. (C) Mean sizes of the total functional group and the strong, positive, recurrent subsets against the population size (i.e. the number of imaged neurons).

### 3.6. The size of the informative functional group saturates

Previously we had found that the accuracy of an encoding model increased with the number of neurons imaged in a dataset (Dechery and MacLean 2018). We set out to establish whether this positive relationship was due to more of each neuron’s respective functional group being sampled, or whether we had imaged a greater number of complete functional groups. As expected the total number of incoming edges increased with the number of neurons imaged per dataset (Fig. 6C – slope=0.21). However, this was not the case when isolating the positive bidirectional edges in the upper quartile of weights. Rather the number of edges in this subset remained relatively stable regardless of the number of neurons imaged (slope was 0.028; Fig. 6C). These results suggest that the increase in model accuracy with increasing numbers of neurons sampled is the consequence of capturing a greater number of complete functional groups. Correspondingly, more neurons are accurately modeled instead of obtaining a better delineation of neurons’ functional groups. This result suggests a numerical size for a functional group.

### 3.7. Functional weights enhance single trial decoding

While the functional group enables accurate single trial encoding predictions, it remained unclear if functional groups are computationally relevant to decoding. More concretely, we asked if using the unrestricted GLM, in which the structure of the functional group is known, to decode the stimulus would result in better performance than decoding when neurons are assumed to be uncoupled since the decoding performance is suggestive of the computational relevance of the functional group (Pillow et al., 2008).

We constructed coupled and uncoupled decoders under the Bayesian decoding framework. In particular, a uniform prior over all stimulus conditions was adopted and the stimulus was decoded via the *maximum a posteriori* estimate. All frames (i.e. both training and test frames) were decoded, and both decoders performed better than chance (Fig. 7A), where chance performance is given by 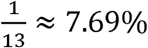 (12 stimulus conditions and 1 grey condition). The coupled GLM decoder decoded more accurately than the uncoupled decoder (coupled GLM mean accuracy = 64.77%, sem = 2.30%, uncoupled mean accuracy = 40.90%, sem = 2.74%, accuracy pooled over all frames, mean over datasets), indicating that knowledge of coupling aids in the extraction of sensory information from the population response. We further computed the mutual information between the true stimulus and the predicted stimulus (Quian Quiroga & Panzeri, 2009), determining the coupled GLM decoder extracts roughly 64% more sensory information (mean = +63.94%, sem = 14.69%, mean over datasets, Fig. 7B). These results indicate, supposing downstream elements have the theoretical capability to read out the visual stimulus, that the functional group dramatically enhances the capability of decoding of the stimulus.

**Fig. 7:**
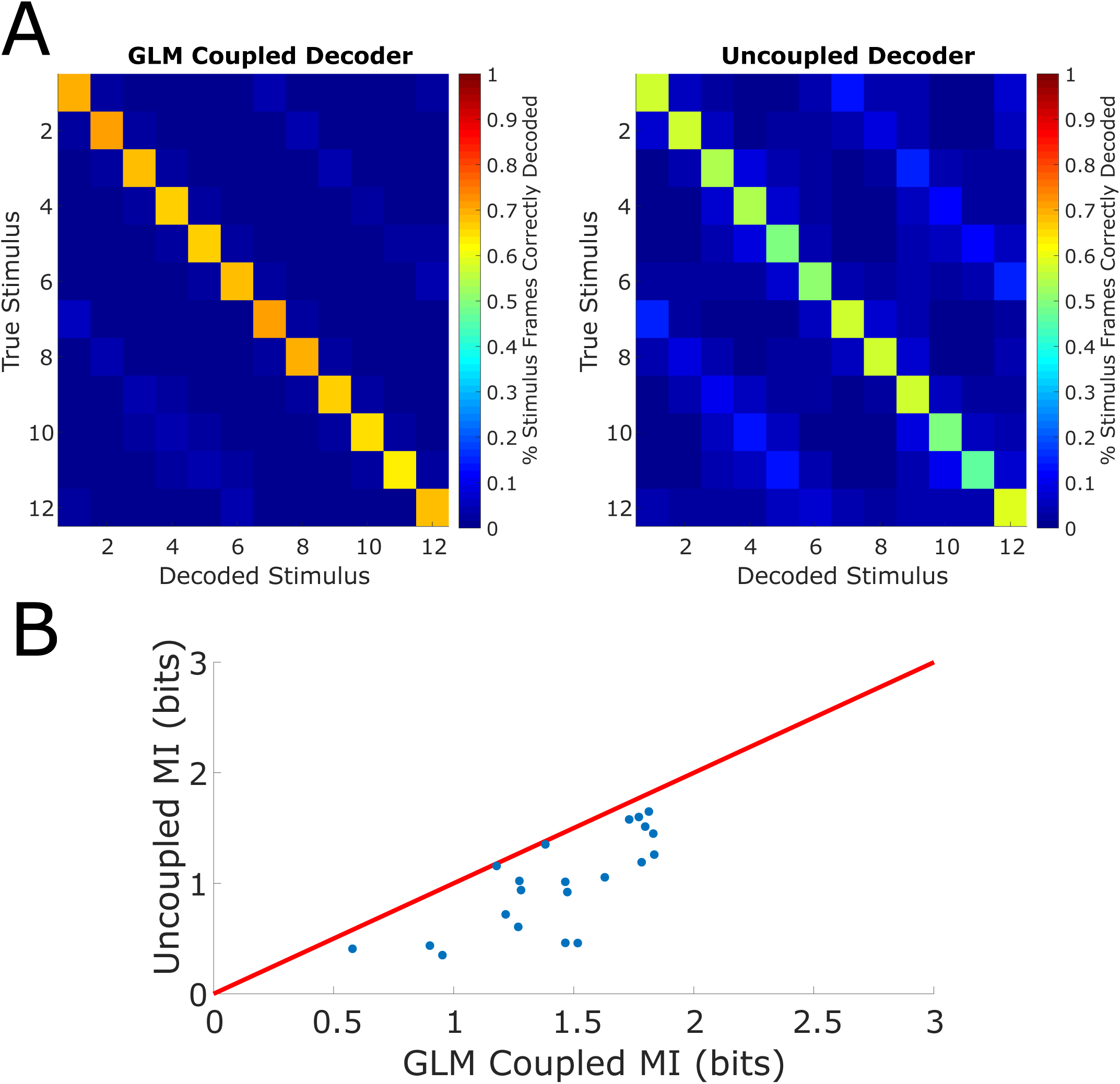
(A) Left, confusion matrix (percent of corresponding stimulus frames correctly decoded) of GLM Coupled Decoder. Right, confusion matrix of Uncoupled Decoder. The GLM Coupled Decoder decodes at a higher accuracy than the Uncoupled Decoder across all stimulus conditions. (B) Comparison of mutual information (MI) between GLM Coupled Decoder and Uncoupled Decoder (red line is unity).

## 4. Discussion

Knowledge of the coactivity of a V1 neuron’s functional group enables accurate predictions of short term dynamics within single trials. Global descriptions such as averaged stimulus-dependent responses and running fail to meaningfully capture single trial activity in the vast majority of L2/3 neurons in visual cortex. With the functional group operationalizing the notion of a Hebbian assembly, our results give concrete descriptions of the timescales, numerical scale, correlation structure, and computational capabilities of assemblies.

### 4.1. Modelling fluorescence with GLMs

The assembly hypothesis posits that internal state dynamics of the underlying circuit shape single trial dynamics; consequently, assembly coactivity is proposed to capture single trials to a greater extent than global variables. To test this proposition, we used various GLMs as encoding models (Paninski, Pillow, & Lewi, 2007; Park, Meister, Huk, & Pillow, 2014; Pillow et al., 2008; Runyan, Piasini, Panzeri, & Harvey, 2017). We used partial correlations to capture shared variability and act as coupling coefficients in the GLM rather than relying on fitted coefficients (Park et al., 2014; Pillow et al., 2008; Runyan et al., 2017). Using empirically measured correlations for coupling coefficients is an attractive modeling choice; fitted coupling coefficients necessarily depend on the model and variable specification, and are thus at risk of omitted variable bias (Stevenson, 2018). Since we use the functional group as an operational notion of an assembly, coupling must not change with specifications of the linear model.

### 4.2. Functional group structure and prediction

We found that the unrestricted GLM accurately modeled the dynamics of single trial activity, thus justifying the decision to adopt a linear model, use partial correlations, and restrict to lags 0 and 1. The target neuron’s variability captured by the model is a deterministic function of the variabilities of the functional group, averaged stimulus-dependent response, and running speed and did not require an explicit stochastic term. Our model results showed functional group coactivity was the main predictor of single trial activity of V1 neurons, in support of the assembly perspective (Harris et al., 2003) as well as previous work showing that pairwise correlations can explain activity patterns (Schneidman et al., 2006; Shlens et al., 2006).

Investigating the robustness of prediction to modulations of the functional group, we found that modulating the topology of large weights severely degrades the model’s ability to predict single trial responses. Hence, the precise instantiation of the strong weights and edges in the functional group is essential for prediction. While global population coupling and fluctuations have been shown to be informative to predicting activity (Clancy, Orsolic, & Mrsic-Flogel, 2019; Lin et al., 2015; Okun et al., 2015; Scholvinck et al., 2015; Stringer et al., 2019), somewhat consistent with our block averaged stimulus and population average response locomotion models, our results point to the inclusion of a local description when predicting neuronal dynamics within single trial responses. Our results largely suggest that strong functional group coupling (a local description) disproportionately captures dynamics, in congruence with the assembly hypothesis and other previous work (Ohiorhenuan et al., 2010).

With the functional group established as the primary predictor of single trials, we investigated its features. Functional group edges with functional lag 0 are sufficient for accurate predictions, suggesting that assembly coactivity largely occurs on timescales of ∼30*ms*, as found in other regions (Harris et al., 2003). Furthermore, while it is known that correlations between neurons are predominantly positive and stronger between similarly tuned neurons (Abbott & Dayan, 1999; Kohn & Smith, 2005; Shadlen & Newsome, 1998), we additionally found that precisely the large, positive weights most enabled accurate predictions.

Using a Bayesian decoding framework, we found that knowledge of the functional improves decoding over treating neurons as independent units, corroborating previous work (Ecker, Berens, Tolias, & Bethge, 2011; Maynard et al., 1999) and suggesting that shared variability is computationally relevant. While the Bayesian decoder does not reveal whether downstream circuits actually have access to or use the functional group structure, these results nonetheless demonstrate the theoretical relevance of functional groups (Quian Quiroga & Panzeri, 2009).

The use of the functional group as an operational definition of an assembly enables concrete descriptions of the timescale, correlation structure, and computational capability of a coactive population. The phenomenon that precise details of the functional group are critical to accurate predictions of dynamics over short time scales indicates that sensory representation and computation are comprised of local coupling in addition to global population-wide variables.

## 5. Methods

### 5.1. Data

A subset of imaging data (n = 21 datasets) of mouse visual cortical neurons was taken from (Dechery & MacLean, 2018). As described in (Dechery & MacLean, 2018), data was collected from n = 4 male and 4 female C57BL/6J mice expressing transgene Tg(Thy1-GCaMP6s)GP4.12Dkim (Jackson Laboratory). Activity of L2/3 excitatory neurons in mouse visual cortex (73–347 neurons; 25–33 Hz; 21 distinct fields of view) were imaged using high speed two photon laser scanning microscopy (Dechery and MacLean 2018; Sadovsky et al 2011). Mice were awake, head-fixed, and allowed to freely run on a linear treadmill while viewing drifting square-wave gratings presented in 12 directions in pseudo-random order interleaved with mean-luminance matched grey screen.

### 5.2. Functional Weights

Partial correlations were used as coupling coefficients in our GLMs; we called these *functional weights*. The functional weight between a pair of neurons is the partial correlation between the corresponding fluorescence traces accounting for the stimulus and population-wide co-activity driven by running. More precisely, for each 5-minute block of oriented drifting grating stimulus presentations (one trial), a trial-specific partial correlation between neuron *x* and neuron *y* was computed while accounting for the mean response of neuron *x* in all other trials, the mean response of neuron *y* in all other trials, and the mean response of the population excluding neurons *x* and *y* in the current trial. Then the trial-specific partial correlations were averaged across trials to obtain the final partial correlation between neuron *x* and neuron *y*; this was taken to be the functional weight between neurons *x* and *y*. The mean activity of neurons *x* and *y* across all other trials was accounted for in order to control for stimulus-dependent effects resulting in a measure analogous to noise correlations (Dechery & MacLean, 2018). The mean population activity excluding neurons *x* and *y* in the current trial is accounted for in order to control for population co-activity driven by running (Dechery & MacLean, 2018). We used the MATLAB 2016a function `parcor.m` to compute partial correlations.

In order to capture the temporal structure of neuron coactivity, functional weights were given directionality in the following manner. The mean fluorescence trace (averaged across trials) for neuron *x* and *y* neuron was computed, and the cross-correlogram was computed. Due to the nature of two-photon imaging, neuronal fluorescence time series are given by a series of time bins with roughly 30 *ms* width; consequently, lags are given in units of time bins. The lag corresponding to the maximum of the cross-correlogram determined the lag of the functional weight and the sign of the lag determined the direction of the functional weight (Dechery & MacLean, 2018), and this lag was termed the *functional lag*. More concretely, if neurons *x* and *y* had a functional weight with a positive functional lag, then the corresponding functional weight has direction corresponding to initial point *x* and terminal point *y*. If the lag is negative, then the functional weight has direction corresponding to initial point *y* and terminal point *x*. Lag 0 corresponds to a bidirectional functional weight. The *functional edge* refers to this directed functional connection. Cross-correlograms were calculated using the MATLAB 2016a functions `xcorr.m`. For each neuron, the set of neurons with outgoing functional edges to the given neuron (i.e. neurons with functional edges with which the given neuron was the terminal neuron) was called the *functional group* of the given neuron.

### 5.3. Generalized Linear Model

#### 5.3.1 Predicting Fluorescence

A generalized linear model (GLM) was used as an encoding model to model calcium imaging data from V1 neurons (Kass, Eden, & Brown, 2014; Park et al., 2014; Pillow et al., 2008; Runyan et al., 2017). The GLM framework allows us to determine which variables are important in predicting the target neuron’s activity by iteratively excluding variables of interest and comparing prediction accuracy.

Stimulus effects were modeled in two ways in order to determine the extent to which averaged summaries of stimulus-dependent responses contribute to accurate predictions of single trial dynamics. First, we modeled the stimulus by the stimulus-dependent average response (which we refer to as “average response”), in which response to a given stimulus was averaged across all time bins of the given stimulus presentation across all trials. Neurons significantly tuned to orientation or direction were labeled as *tuned* with the procedure described in (Dechery & MacLean, 2018), and all remaining neurons were labeled as *untuned*. The stimulus term in our GLM was the average stimulus-dependent response for tuned neurons. The stimulus term for untuned neurons was given by the response given by averaging the stimulus-dependent average response across all stimulus conditions. During grey frames, the stimulus term was set to zero in this model. With this model, we used a 70/30 split of the data for training and testing in order to fit the GLM coefficients and test performance. We modeled the stimulus a second way by the “block averaged trace” in order to reflect dynamics associated to neuronal response. In this model, tuned and untuned neurons were treated exactly the same. The fluorescence traces in each presentation of a given stimulus were averaged across blocks (i.e. trials) but not time bins, producing a block-averaged trace which preserved the dynamics. Since the block-averaged trace preserves dynamics, we only take averages over the training set when fitting the GLM to avoid overfitting (when testing, averages are taken over the testing set). In order to ensure that averages occurred over similar numbers of stimulus presentations between the training and testing sets, we used a 50/50 split of the data. In our analyses, the block averaged trace was obtained after averaging over 9-15 traces depending on the dataset. The stimulus term is represented by *s*(*t*) in our GLM equation. Running effects were modeled without neuron-specific modulation, i.e. we included a term *v*(*t*) (denoting the running velocity at time *t*) that was constant across all neurons.

The functional group was incorporated into the GLM by computing a linear combination (with weights given by the functional weights) of the lagged (lag given by the functional lag) fluorescence traces of all neurons in the functional group (i.e. neurons with outgoing functional edges to the given neuron). For compact notation, we let ***r***(*t*) be the vector of population fluorescence at a given time, *T* be the total duration of the event, and *N* be the total number of neurons in the population. Then we defined 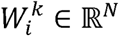 to be the vector of functional weights corresponding to incoming edges to neuron *i* with lag *k*. In the entire investigation, we restricted attention to *k* ∈ {0,1} as the frequency of lagged functional relationships with *k* > 1 decreased rapidly.

With the coupling, stimulus term, and running term defined, for each neuron *i*, we constructed a GLM to model its fluorescence,

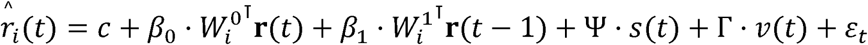

Here, *c, β*_0_, *β*_1_, Ψ, Γ ∈ ℝ are coefficients and *ε*_*t*_ is mean zero noise. The linear offset *c* is needed as different neurons have different baseline fluorescence. The gain terms *β*_0_, *β*_1_ are needed to account for the range of incoming edges to each neuron. The Ψ and Γ coefficients are the modulation associated with stimulus and running respectively. The notation 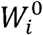 and 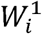 refers to the vector of lag 0 and lag 1 incoming functional weights to neuron *i*. More specifically, 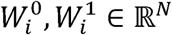, and

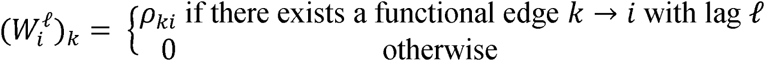

Here, *ρ*_*ki*_ is the functional weight associated to the directed edge *k* → *i*. Since the vectors of functional weights 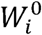 and 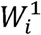 are computed directly from the data, the scalars 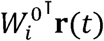 and 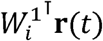 are features of the data and *not* free parameters. Hence, for a given neuron, there exist 5 parameters in the unrestricted GLM. For each neuron, the model was fit on the training data by the method of least squares using a MATLAB R2016a function (`lscov.m)`.

### 5.4. GLM Variants

#### 5.4.1 Stimulus-restricted, Run-restricted, and Coupling-restricted GLMs

To determine the extent to which the stimulus term, running term, and coupling terms contributed to predictions of single trials, we iteratively excluded terms of interest, refit the model, and compared predictive performance against the unrestricted model (i.e. containing all model terms described above). In particular, for the *stimulus-restricted GLM*, we had the model

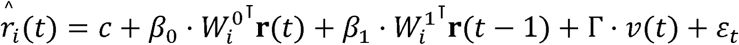

in which the stimulus term was excluded. The model coefficients all have the same interpretation as described in the previous section. We then fit this model on the training data, and compared its predictive performance on the test data with the unrestricted model. Similarly, for the *run-restricted GLM*, we had the model

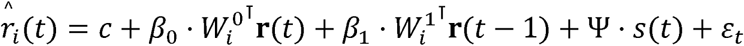

and for the *coupling-restricted GLM*, we had the model

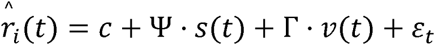

These models were fit on the training data, and were compared against the unrestricted model in terms of predictive performance on the test data.

#### 5.4.2 GLM_0_ and GLM_1_

To determine the contributions of lag 0 and lag 1 coupling to predictions of single trials, we similarly excluded these terms iteratively and compared predictive performance to the unrestricted model. In particular, *GLM*_*0*_ is given by

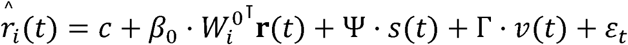

This model was then fit on the training data and compared against the unrestricted model. Note that 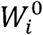 is not affected by refitting as it determined by the functional weights and functional lags, which are empirically measured. The process of fitting only affects the model coefficients, i.e. *c, β*_0_, Ψ, Γ. Similarly, *GLM*_*1*_ is given by

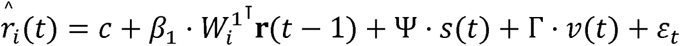

and the model is fitted and evaluated with exactly the same procedure as *GLM*_*0*_.

#### 5.4.3 GLM^+^ and GLM^-^

To determine the contributions of positive and negative functional weights to predictions of single trial dynamics, we constructed a GLM with explicit corresponding terms (again, with average response as the stimulus term). In particular, we had the following model

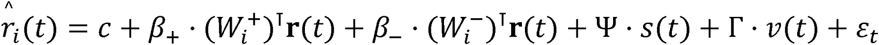

The terms **r**(*t*), *s*(*t*), *v*(*t*) and *ε*_*t*_ as in the original model formulation. The notation 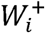 and 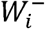 refers respectively to the positive and negative functional weights in the functional group of neuron *i*. More specifically, 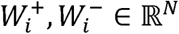, and for neurons *k* and *i* and sign *s* (i.e. positive or negative), we had

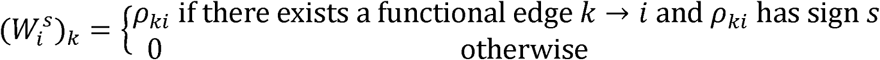

In order to minimize over-fitting and keep model complexity comparable across different models, we used all lags in this model and treated them as lag 0.

### 5.5. Permutations of Functional Group Topology

To determine if the topology of the function group is essential to predictions of single trial activity, we shuffled the functional group in two ways; we shuffled the “strong weights” and the “strong edges”.

We shuffled the strong weights to determine the importance of strong weights for prediction. We shuffled the top quartile of weights while preserving the corresponding underlying edges. Formally, we enumerated the edges with strong weights in our entire population {(*i*_1_, *j*_1_, *w*_1_),…,(*i*_*K*_, *j* _*K*_, *w* _*K*_)}, where a directed edge from neuron *i* _*α*_ to neuron *j* _*α*_ exists and *w* _*α*_ is the associated weight for 1 ≤ *α* ≤ *K*. This enumeration contains all tuples (*i* _*α*_, *j* _*α*_, *w* _*α*_, such that |*w* _*α*_| is in the top quartile of all weight magnitudes. We then applied a uniformly random permutation *σ* ∈ *S*_*K*_ (where *S*_*K*_ is the finite symmetric group on *K* points) to act on the weights, furnishing the enumeration {(*i*_1_, *j*_1_, *w*_*σ*(1)_),…,(*i*_*K*_, *j* _*K*_, *w*_*σ*(*K*)_)}. We then assigned these permuted weights to the corresponding edges (i.e. the directed edge from neuron *i* _*α*_ to neuron *j* _*α*_ was assigned the weight *w*_*σ*(*α*)_) in the functional group. Furthermore, we shuffled the functional lags in this way (thus changing the directionality of the underlying edges). This construction preserves the existence of edges, preserves which edges are in the top quartile of weight magnitudes, and preserves structure between functional lags and functional weights. Hence, changes in prediction accuracy are directly related to the importance of the functional weight to prediction.

Similarly, to isolate the importance of the functional edges in prediction, we shuffled the strong edges of the functional group. We apply a uniformly random permutation *σ* ∈ *S*_*K*_to the list of terminal neurons. Formally, with the list {(*i*_1_, *j*_1_, *w*_1_),…,(*i*_*K*_, *j* _*K*_, *w*_*K*_)} described above, we apply the permutation to obtain {(*i*_1_, *j*_(1)_, *w*_*σ*(1)_),…,(*i*_*K*_, *j* _*K*_, *w*_*σ*(*K*)_)}. This procedure permutes the strong edges while preserving the weight with respect to the source neuron. Hence, changes in prediction accuracy are directly related to the importance of functional edges to prediction.

### 5.6. Decoding

#### 5.6.1 Constructing the decoder

To determine whether the functional group is computationally relevant, we used the functional group and GLM framework to decode the stimulus. We constructed a Bayesian decoder which estimates the stimulus at time *t* given the population response and the running speed at time *t*. The unrestricted GLM (using average response for the stimulus term) was incorporated into the decoder (Paninski et al., 2007) in order to directly link the functional structure of the population to decoding; this was termed the coupled decoder. The coupled decoder was then compared to an uncoupled decoder: a Bayesian decoder which treats all units as conditionally independent (Pillow et al., 2008; Runyan et al., 2017).

The coupled decoder was constructed as follows. From Bayes’ Theorem, the posterior probability of stimulus conditional on response is proportional to the product of the prior and the probability of response conditional on stimulus. Formally,

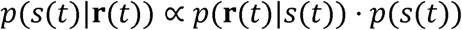

Here, **r**(*t*) is the vector of population activity and *s*(*t*) is the stimulus at time *t*. Estimating *p*(*s*(*t*)|**r**(*t*)) requires specification of *p*(*s*(*t*)) and *p*(**r**(*t*)|*s*(*t*)), which we gave as

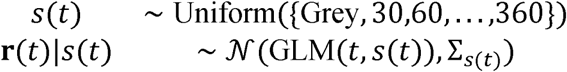

Here, GLM (*t,s*(*t*)) is the vector of GLM predicted responses at time *t* conditioned on the stimulus input *s*(*t*). Further, Σ_*s*(t)_ is the covariance of the neurons’ traces during frames with *s*(*t*) presentation. The decoded stimulus is then the *maximum a posteriori* estimate (MAP)

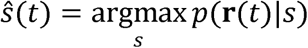

This is standard in Bayesian decoders (Paninski et al., 2007; Quian Quiroga & Panzeri, 2009). The uncoupled decoder is constructed similarly. The prior and conditional distributions are specified as

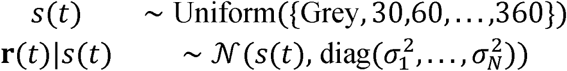

Here, *s*(*t*) = (*s*_1_(*t*),…,*s*_*N*_(*t*)) is the vector of stimulus-dependent average response. Further, 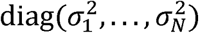 is the diagonal matrix with 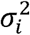 denoting the variance of neuron *i*’s fluorescence trace during all frames when the stimulus *s*(*t*) was presented. Note that since the conditional distribution posits a diagonal covariance matrix in a multivariate Gaussian, the neurons are independent, and we have the factorization

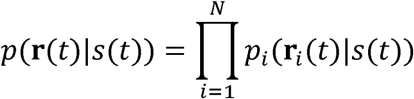

Here, *p*_*i*_ is the marginal density of neuron *i*’s trace. Similarly, the decoded stimulus is the MAP estimate. Using the 70/30 data split for training and testing from previous analyses, we fit the GLM on the training set, then decoded all of the frames (both training and testing frames) in our analysis.

#### 5.6.2 Estimation of information about stimulus

The extent to which sensory information was extracted from the population response was investigated via an information theoretic approach. Specifically, we computed the mutual information between the decoded stimulus and the true stimulus (Quian Quiroga & Panzeri, 2009), given by

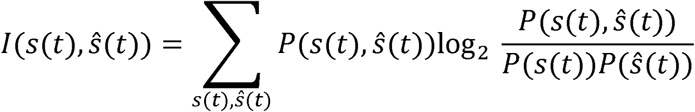

Here, *s*(*t*) is the true stimulus, *ŝ*(*t*) is the decoded stimulus, and the mutual information *I*(*s*(*t*),*ŝ*(*t*)) has units in bits. The joint and marginal distributions were estimated directly from the decoding results. Given the large size of our data, subsampling the distributions was not a concern. This approach gave a meaningful quantification of information and enabled sensible comparisons between decoders. Further, it gave a measure for the computational relevance of the functional group beyond single trial predictive power.

## Acknowledgments

This work was supported by NIH grant R01EY022338. The authors wholeheartedly thank Joseph Dechery for collecting the mouse data sets used in the analysis. Additionally, the authors thank John Maunsell, Jackson Cone, Maayan Levy, and Elizabeth de Laittre for their helpful and deliberate comments on the manuscript.

